# Escalation of genome defense capacity enables control of an expanding meiotic driver

**DOI:** 10.1101/2024.06.12.598716

**Authors:** Peiwei Chen, Katherine C. Pan, Eunice H. Park, Yicheng Luo, Yuh Chwen G. Lee, Alexei A. Aravin

**Affiliations:** Division of Biology and Biological Engineering, California Institute of Technology, Pasadena, California 91125, USA; Department of Ecology and Evolutionary Biology, University of California, Irvine, Irvine, California 92697, USA

## Abstract

**Summary:** From RNA interference to chromatin silencing, diverse genome defense pathways silence selfish genetic elements to safeguard genome integrity^1,2^. Despite their diversity, different defense pathways share a modular organization, where numerous specificity factors identify diverse targets and common effectors silence them. In the PIWI-interacting RNA (piRNA) pathway, which controls selfish elements in the metazoan germline, diverse target RNAs are first identified by complementary base pairing with piRNAs and then silenced by PIWI-clade nucleases via enzymatic cleavage^1,3^. Such a binary architecture allows the defense systems to be readily adaptable, where new targets can be captured via the innovation of new specificity factors^4,5^. Thus, our current understanding of genome defense against lineage-specific selfish genes has been largely limited to the evolution of specificity factors, while it remains poorly understood whether other types of innovations are required. Here, we describe a new type of innovation, which escalates the defense capacity of the piRNA pathway to control a recently expanded selfish gene in *Drosophila melanogaster*. Through an *in vivo* RNAi screen for repressors of *Stellate*—a recently evolved and expanded selfish meiotic driver^6–8^—we discovered a novel defense factor, Trailblazer. Trailblazer is a transcription factor that promotes the expression of two PIWI-clade nucleases, Aub and AGO3, to match *Stellate* in abundance. Recent innovation in the DNA-binding domain of Trailblazer enabled it to drastically elevate Aub and AGO3 expression in the *D. melanogaster* lineage, thereby escalating the silencing capacity of the piRNA pathway to control expanded *Stellate* and safeguard fertility. As copy-number expansion is a recurrent feature of diverse selfish genes across the tree of life^9–12^, we envision that augmenting the defense capacity to quantitatively match selfish genes is likely a repeatedly employed defense strategy in evolution.

## Main Text

The genome is a battleground, where every bit of DNA competes for inheritance to the next generation^13^. In particular, selfish genes enhance their own propagation at the expense of the host fitness, causing intragenomic conflicts that must be resolved to protect host reproduction^14,15^. To keep selfish genes in check, organisms employ diverse genome defense mechanisms, such as RNA interference (RNAi) and chromatin silencing guided by Krüppel-associated box domain zinc finger proteins (KRAB-ZFPs)^1,2^. Despite their diversity, different genome defense mechanisms often share a binary core architecture: a specificity factor identifies the target, and an effector confers its silencing. For example, in small RNA silencing pathways, a short RNA finds the target RNA based on complementary base pairing, while the associated Argonaute protein silences the target, *e*.*g*., by cleaving its RNA^1,3^. Such a binary core architecture enables the adaptability of defense systems: by innovating a new specificity factor, defense pathways can be directed against a new target, without changes in effectors or other components. Indeed, genetic innovations are repeatedly found in specificity factors of diverse defense pathways across the tree of life, paralleling the recurrent emergence of distinct selfish genes in different lineages^4,5^. Thus, the current view of how lineage-specific selfish genes are controlled is largely limited to innovations of specificity factors, such as the evolution of new small RNAs or new KRAB-ZFPs. Meanwhile, it remains largely unexplored whether genome defense against evolutionarily young selfish genes requires other types of genetic innovations outside specificity factors.

To decipher how evolutionarily young selfish genes are tamed, we focused on a recently evolved selfish gene family, *Stellate*, which is only active in the male germline of *Drosophila melanogaster. Stellate* is a multi-copy gene family on the X chromosome ^16^ (Fig. 1A), and it is thought to be a meiotic driver that promotes the inheritance of X-bearing sperm at the cost of Y-bearing counterparts^6–8^. Indeed, de-repression of *Stellate* in the male germline causes meiotic and post-meiotic defects^16–18^ that result in either female-biased offspring sex ratios (when *Stellate* copy number is low) or complete sterility (when *Stellate* copy number is high)^6^. The defects caused by Stellate are suppressed by the *Suppressor of Stellate* [*Su(Ste)*] locus on the Y chromosome, which encodes PIWI-interacting RNAs (piRNAs) that direct two PIWI-clade Argonaute proteins, Aubergine (Aub) and Argonaute 3 (AGO3), to cleave *Stellate* mRNAs^19–23^ (Fig. 1A). In other words, the innovation of specificity factors, *Su(Ste)* piRNAs, allowed the conserved piRNA pathway machinery to recognize and repress recently evolved *Stellate*. Whether *Stellate* silencing requires other genetic innovations beyond *Su(Ste)* piRNAs, however, remained unknown. Of note, silencing *Stellate* is critical for male fertility, while suppression of other piRNA targets (including diverse transposons) appears less important. Specifically, de-repression of *Stellate* alone can render males sterile^6^, whereas disrupting global piRNA biogenesis without perturbing *Su(Ste)* piRNAs does not lead to male infertility^24^. The indispensability of *Stellate* silencing, therefore, makes it an attractive model to explore whether developing robust genome defense requires innovations beyond target-specific piRNAs.

**Fig. 1.**
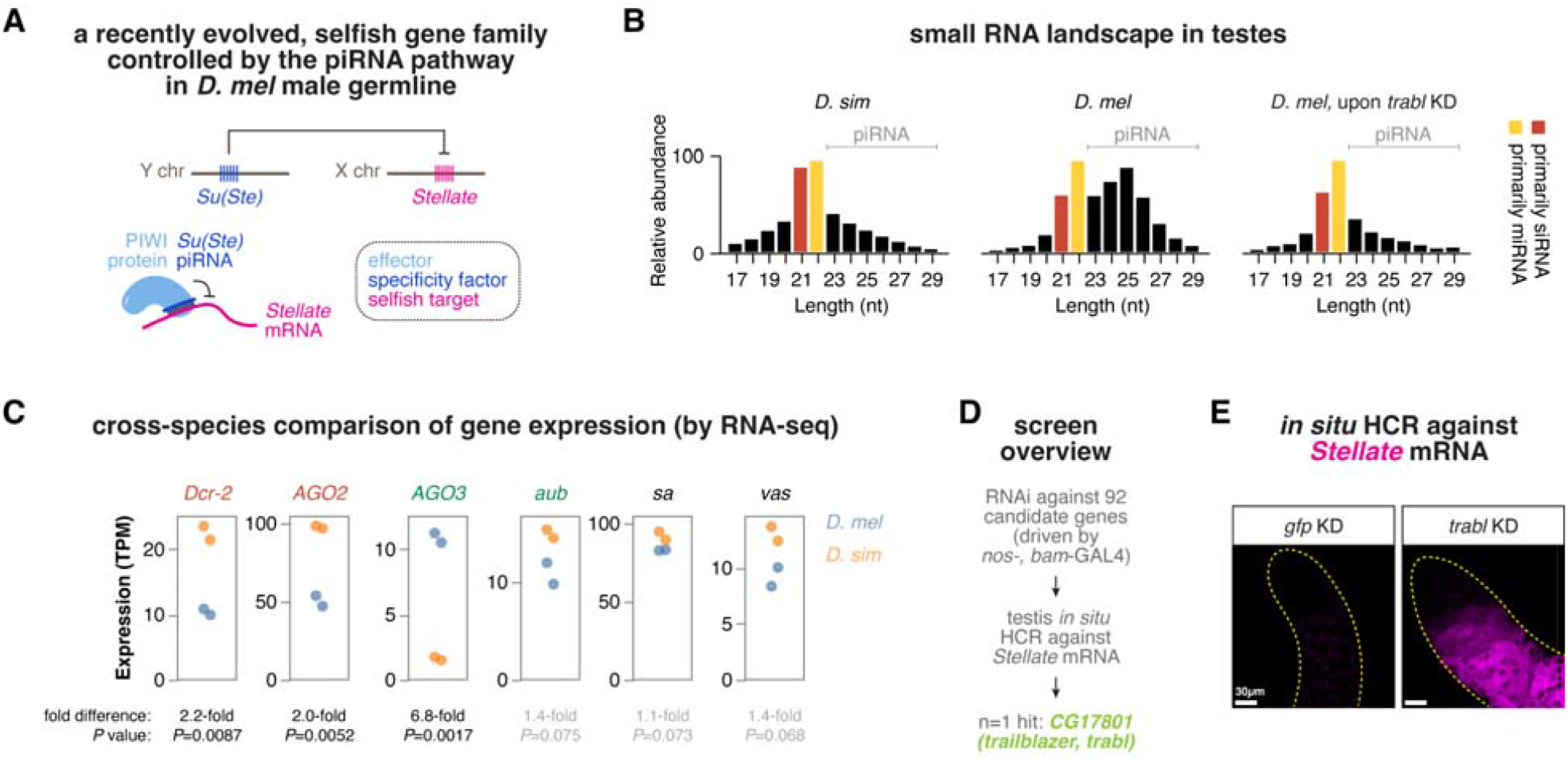
Control of the selfish *Stellate* genes in *D. melanogaster* is accompanied by radical changes in the genome-defending piRNA pathway. (**A**) Schematic showing the recently evolved, X-linked selfish *Stellate* gene family controlled by Y-linked *Su(Ste)* piRNAs in *D. melanogaster. Stellate* and *Su(Ste)* are absent in *D. simulans* and other closely related species. In the genome-defending piRNA pathway, piRNAs are specificity factors that identify selfish targets, while PIWI proteins are effectors that confer silencing. (**B**) Small RNA profiles in testes of *D. simulans, D. melanogaster*, and *D. melanogaster* upon *trabl* germline KD. 21nt and 22nt small RNAs that are primarily siRNAs and miRNAs are marked red and yellow, respectively. (**C**) mRNA expression of *Dcr-2, AGO2* (core factors of the siRNA pathway), *AGO3, aub* (core factors of the piRNA pathway) and *sa, vas* (germline markers) in *D. melanogaster* (blue) and *D. simulans* (orange) testes. P-values from unpaired t-tests are shown along with fold differences between species. (**D**) An RNAi screen identified a novel genome defense factor, Trailblazer. Overview of the RNAi screen for repressors of *Stellate*. (**E**) RNA *in situ* HCR against *Stellate* mRNAs in testes upon *trabl or* control (*gfp*) germline KD (driven by *nos-* and *bam-*GAL4).

Analysis of small RNAs in testes of *D. melanogaster* and its closely related species, *Drosophila simulans*, which lacks *Stellate* and *Su(Ste)*, supports the idea that the evolution of *Stellate* repression is associated with radical changes in genome defense. When compared to small interfering RNAs (siRNAs) and microRNAs (miRNAs), piRNAs as a whole are of similar abundance as other small RNAs in *D. simulans* testes (piRNA-to-miRNA ratio is about 0.91; Fig. 1B). In contrast, piRNAs are exceptionally abundant in *D. melanogaster* testes, with 2.81 piRNA-to-miRNA ratio. Furthermore, nearly half of the total piRNA pool in *D. melanogaster* testes is occupied by *Su(Ste)* piRNAs, outnumbering all transposon-targeting piRNAs combined^22,25^. These observations suggest that the control of recently evolved *Stellate* genes in *D. melanogaster* is associated with an augmentation of the overall piRNA pathway activity and a quantitative increase in the piRNA population.

To investigate if the exceptional abundance of piRNAs in *D. melanogaster* testes relative to *D. simulans* testes is accompanied by other cross-species differences, we analyzed their gene expression profiles. Testis transcriptome profiling by RNA-seq showed that the expression of *Argonaute 2 (AGO2)* and *Dicer-2 (Dcr-2)*—two key components of the siRNA pathway—is 2-fold higher in *D. simulans* than *D. melanogaster* (p-values<0.01) (Fig. 1C), consistent with the higher relative abundance of siRNAs in *D. simulans* than *D. melanogaster* (Fig. 1B). In contrast, our examinations of the expression of *aub* and *AGO3*, which encode for the two PIWI proteins that cleave *Stellate* mRNAs, gave a strikingly different picture. *aub* expression is comparable between the two species, and even more, the expression of *AGO3* is 7-fold higher in *D. melanogaster* than *D. simulans* (p-value: 0.0017) (Fig. 1C). The elevated *AGO3* expression in *D. melanogaster* testes is unlikely to be driven by a higher fraction of germ cells or a general increase of germline gene expression, because the expression of *vasa* and *spermatocyte arrest (sa)*—two germline genes that are co-expressed at the same stages of spermatogenesis as *aub/AGO3* and *Stellate*, respectively—is slightly higher in *D. simulans* (Fig. 1C). Together, our analysis of small RNAs and gene expression profiles suggests that the evolution of *Stellate* repression by the piRNA pathway in *D. melanogaster* is associated with significant increases in both the overall piRNA abundance and the expression of a key effector, AGO3.

### An RNAi screen identified a novel genome defense factor, Trailblazer

In addition to Aub and AGO3 that load piRNAs and act as nucleases to degrade complementary RNAs, the piRNA pathway engages more than 30 proteins. Notably, almost all of these proteins were initially identified in females, and they function either in both sexes or in only females^26–29^. This female-biased view might cause the omission of factors critical for males, especially those involved in *Stellate* silencing specifically. To identify genes required for *Stellate* silencing, we conducted a targeted *in vivo* RNAi screen, using *in situ* hybridization chain reaction (HCR) to visualize *Stellate* mRNAs^30^ (Fig. 1D). Our screen focused on the zinc-finger associated domain (ZAD) zinc finger protein family, which drastically expanded in the insect lineage and has 92 paralogs in *D. melanogaster*^31,32^. The expansion of ZAD zinc finger protein family in insects parallels the expansion of KRAB-ZFPs, another zinc finger protein family in tetrapods^33^. The expansion of KRAB-ZFPs has been suggested to be driven by an arms race with the expanding, lineage-specific retrotransposons^4,9^. It is also worth noting that one ZAD zinc finger protein was recently identified to be involved in the piRNA pathway in the *D. melanogaster* female germline^29^. We thus hypothesized that the expanded ZAD zinc finger protein family might include innovations in the defense against *Stellate*.

Out of the 92 ZAD zinc finger genes, our screen identified one gene, *CG17801*, that is required for *Stellate* silencing. Knocking-down (KD) *CG17801* in the male germline using two different RNAi lines led to *Stellate* de-repression (Fig. 1E and fig. S1). Similarly, *CG17801* mutant males harboring two different allele combinations exhibited *Stellate* de-silencing. With a high *Stellate* copy number (see below), males lacking *CG17801* (upon either KD or mutations) were completely sterile (Fig. 2, A and B). Remarkably, such male infertility can be rescued by simultaneously depleting *Stellate* (Fig. 2C), indicating that *Stellate* de-repression is the primary cause of male infertility upon *CG17801* KD. Indeed, males lacking *CG17801* were spermless, but the sperm production was recovered when *Stellate* was depleted concomitantly (Fig. 2D). As opposed to males, female fertility remained normal without *CG17801* (Fig. 2, A and B). Together, these results suggest that *CG17801* encodes a new genome defense factor that represses *Stellate* in males. We named CG17801 Trailblazer, because it is a “pioneer” that tames recently evolved selfish genes.

**Fig. 2.**
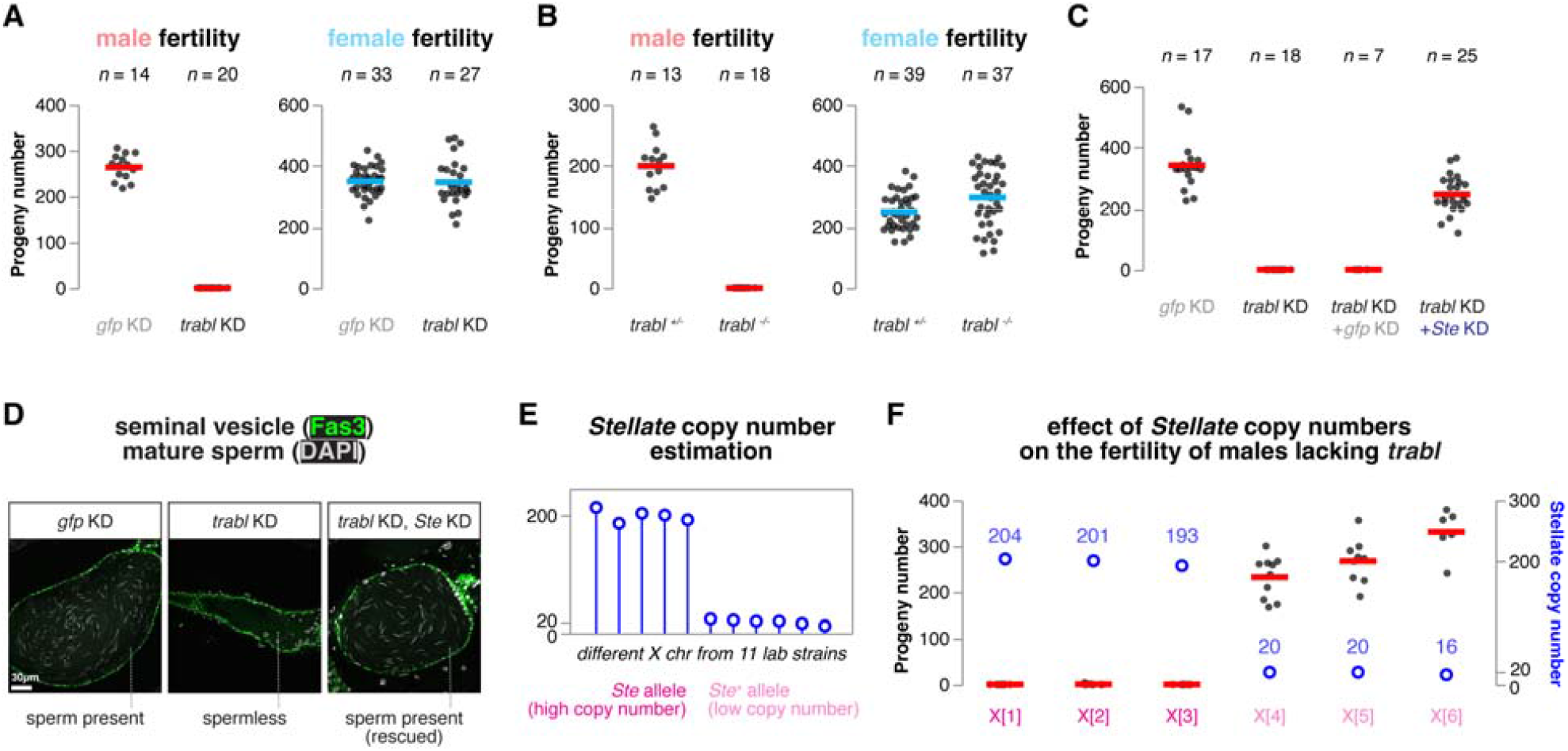
A novel genome defense factor, Trailblazer, silences *Stellate* to safeguard male fertility. (**A**) Fertility tests of males and females upon *trabl* or control (*gfp*) germline KD. Horizontal lines denote the mean. (**B**) Fertility tests of *trabl* heterozygous controls and trans-heterozygous mutants of both sexes. (**C**) Male fertility tests upon *gfp* germline KD, *trabl* germline KD, and *trabl* germline KD with simultaneous *gfp* or *Stellate* KD. (**D**) Immunofluorescence images of seminal vesicles (SVs) stained by Fas3, with needle-shaped sperm head inside SVs marked by DAPI staining. (**E**) Estimation of *Stellate* copy numbers on X chromosomes from 11 lab strains using quantitative PCR. (**F**) Fertility tests of *trabl* mutant male with different *Stellate* copy numbers. Horizontal lines denote mean progeny numbers, while open blue circles mark estimated *Stellate* copy numbers.

The effect of *Stellate* de-repression on male fertility is known to correlate with *Stellate* copy numbers, which vary across different *D. melanogaster* strains^6,16^. Historically, two *Stellate* alleles have been described: “*Ste*”, which has a high copy number and causes male infertility when de-silenced, and “*Ste*^*+*^”, which has a low copy number and does not abolish male fertility when de-silenced^6,16^. We quantified *Stellate* copy numbers in different X chromosomes from 11 lab strains and found that they follow a bimodal distribution: either ∼20 or ∼200 copies, akin to *Ste*^*+*^ and *Ste* alleles described in literature (Fig. 2E). To investigate if lab strains with different *Stellate* copy numbers show differential dependence on Trailblazer, we examined male fertility of *trailblazer* mutants that have different X chromosomes: three with ∼20 *Stellate* genes and another three with ∼200. High *Stellate* copy numbers render mutant males sterile, while low copy numbers of *Stellate* do not (Fig. 2F), indicating that the penetrance of *trailblazer* phenotypes correlates with the copy-number expansion of selfish *Stellate* genes.

### Trailblazer is a transcription factor for *aub* and *AGO3*

We next sought to address how Trailblazer modulates *Stellate* silencing. Trailblazer is only expressed in gonads^34^, and, in testes, its mRNAs are only detected in the early male germline (Fig. 3A). When we examined *Su(Ste)* piRNA precursor transcription, we found that the *Su(Ste)* transcript level remains unperturbed upon *trailblazer* KD (Fig. 3A), arguing against a role of Trailblazer in promoting *Su(Ste)* piRNA precursor transcription. To gain further insights into Trailblazer functions, we performed RNA-seq in testes of males lacking *trailblazer*. Depleting *trailblazer* caused limited changes in the transcriptome: only about 30 genes were differentially expressed, including strongly de-repressed *Stellate* (Fig. 3B). Surprisingly, two genes that are down-regulated upon *trailblazer* KD are *aub* and *AGO3* (Fig. 3, B and C), which encode the two PIWI proteins that act as endonucleases to cleave piRNA targets, including *Stellate*^22,23^.

**Fig. 3.**
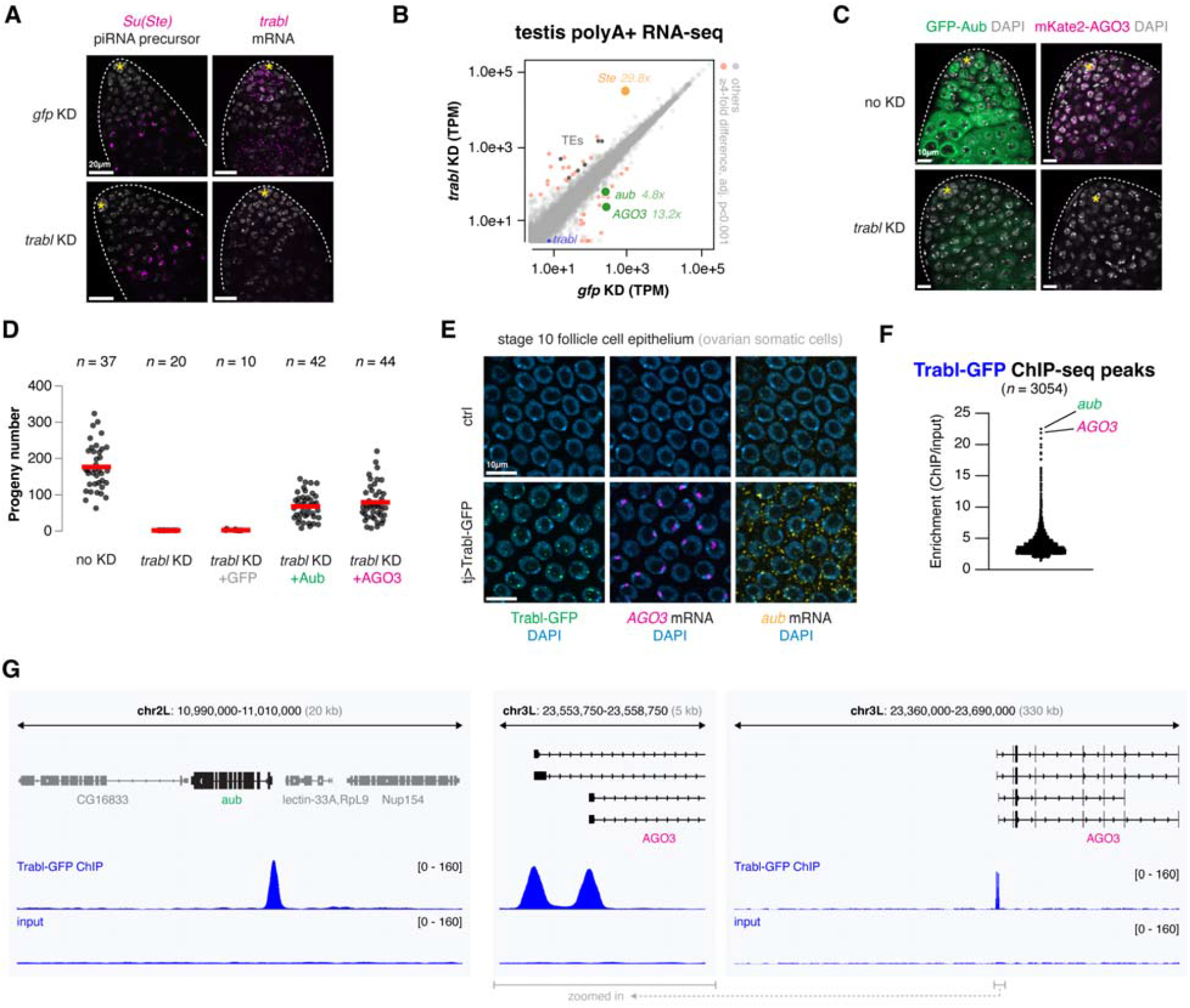
Trailblazer is a transcription factor that promotes *aub* and *AGO3* expression in the germline. (**A**) RNA *in situ* HCR against *Su(Ste)* and *trabl* in testes upon *trabl* germline KD. Yellow asterisks denote the hub. (**B**) Scatter plot showing the differential gene expression in testes upon *trabl* germline KD. (**C**) Confocal images of testes with endogenous, fluorescently tagged Aub and AGO3 upon *trabl* germline KD. (**D**) Male fertility tests upon *trabl* germline KD or *trabl* germline KD with simultaneous GFP, Aub, or AGO3 expression driven by *nos-* and *bam-* GAL4. (**E**) RNA *in situ* HCR against *aub* and *AGO3* in follicle cell epithelium with or without ectopic expression of Trabl-GFP by *tj*-GAL4. Note that nucleoli appear as DAPI-weak circle inside the follicle cell nuclei. (**F**) Violin plot of ChIP enrichment of Trabl genome-wide binding sites. (**G**) Genome browser views of *aub* and *AGO3* promoters being bound by Trabl. Two alternative promoters of *AGO3* are both bound by Trabl.

Unlike the *Su(Ste)* piRNAs that specifically recognize *Stellate* mRNAs, Aub and AGO3 are general effectors that silence the targets identified by their piRNA guides. The lack of target specificity in Aub and AGO3 raises the question of why *Stellate* silencing may be particularly sensitive to their loss. Our transcriptome profiling showed that the abundance of *Stellate* transcripts exceeds that of all transposons by an order of magnitude in testes (Fig. 3B), suggesting that *Stellate* silencing requires a large amount of Aub and AGO3. In agreement with this, we found that male infertility upon *trailblazer* KD can be rescued by simultaneously expressing Aub or AGO3 in the male germline (Fig. 3D). These observations demonstrate that Trailblazer facilitates *Stellate* silencing by promoting the expression of Aub and AGO3—two general effectors in the piRNA pathway.

To elucidate how Trailblazer promotes *aub* and *AGO3* expression, we generated transgenic flies that express GFP-tagged Trailblazer under the control of GAL4/UAS. The *trailblazer* transgene with a C-terminal GFP tag completely rescued the infertility of males lacking endogenous *trailblazer* (Fig. S2), indicating that it is fully functional. When we used *tj*-GAL4 to drive *trailblazer* expression in follicle cells—ovarian somatic cells that do not normally express *trailblazer, aub*, or *AGO3*—transcriptions of both *aub* and *AGO3* were activated (Fig. 3E). This indicates that expressing *trailblazer* in the soma is sufficient to induce the expression of two germline-specific piRNA pathway effectors. When expressed, Trailblazer localizes to the nucleus and forms distinct foci on the chromatin (Fig. 3E), suggesting that it might be involved in transcriptional regulation.

To understand the Trailblazer function in the nucleus, we profiled the genome-wide binding of Trailblazer to chromatin by performing chromatin immunoprecipitation followed by sequencing (ChIP-seq). This was done in ovaries expressing *trailblazer-gfp* throughout germline development, as *trailblazer* is only expressed in very few early germ cells in testes (Fig. 3A) and existing GAL4 lines cannot drive broader expression in testes. ChIP-seq revealed that Trailblazer binds to ∼3,000 distinct sites throughout the genome (Fig. 3F). While most of these sites have relatively weak ChIP enrichment (<5-fold), several sites located at gene promoters showed strong Trailblazer bindings (>15-fold ChIP enrichment). Strikingly, the two strongest Trailblazer binding sites are located at the promoters of *aub* and *AGO3*, including two alternative promoters of *AGO3* (Fig. 3G). Together with the ability of Trailblazer to induce *aub* and *AGO3* expression in the soma, these ChIP-seq results indicate that Trailblazer is a transcription factor that promotes *aub* and *AGO3* expression.

### DNA binding innovation endows Trailblazer the ability to up-regulate *aub* and *AGO3*

Trailblazer is found in many species of the *Drosophila* genus^35^, indicating that it is much older than the recently evolved, *D. melanogaster*-specific *Stellate*. Accordingly, we predict that there must be genetic innovations within Trailblazer to evolve novel functions to combat *Stellate*. We tested this idea by investigating whether the function of *D. melanogaster trailblazer* can be substituted by the ortholog from a closely related species, *D. simulans* (Fig. 4A). Using CRISPR/Cas9, we replaced the endogenous *trailblazer* coding sequence in *D. melanogaster* with the ortholog from *D. simulans*. Despite sharing 82% amino acid sequence identity, *D. simulans trailblazer* is unable to complement the function of its *D. melanogaster* homolog. *D. melanogaster* males expressing Trailblazer^simulans^ failed to represses *Stellate* (Fig. 4B), had reduced *aub* and *AGO3* levels (Fig. 4C), and exhibited severe fertility defects (Fig. 4D). Importantly, such fertility defects can be rescued by depleting *Stellate*, expressing *aub*, or expressing *AGO3* (Fig. 4D), indicating that fertility defects of Trailblazer^simulans^ result from its inability to drive sufficient *aub* and *AGO3* expression for *Stellate* silencing. Therefore, *D. melanogaster*-specific Trailblazer residues are required to activate sufficient *aub* and *AGO3* to tame *D. melanogaster*-specific *Stellate*.

**Fig. 4.**
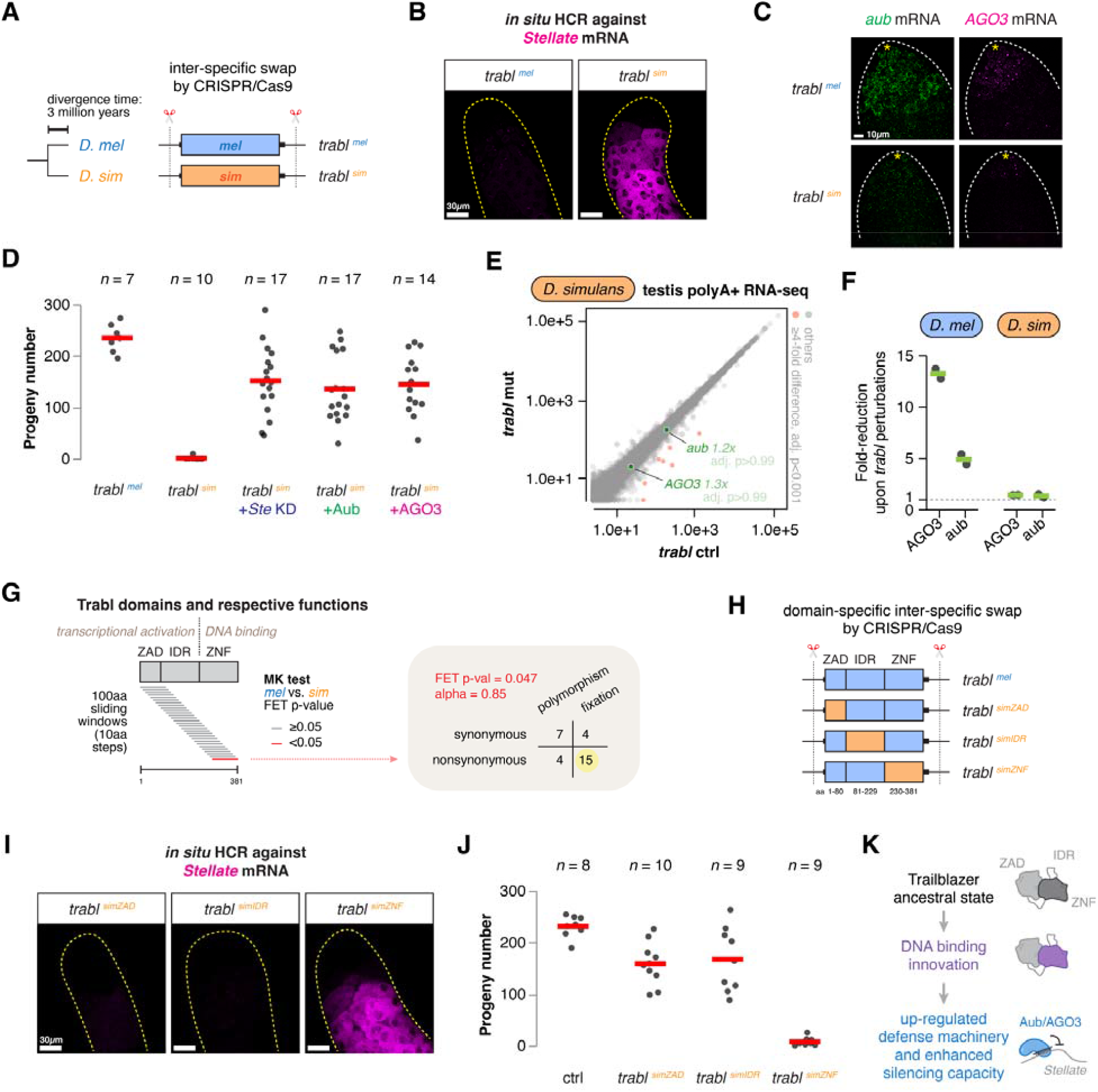
DNA binding innovation endows Trailblazer the ability to up-regulate *aub* and *AGO3*. (**A**) (left) Divergence of *D. melanogaster* from its sibling species *D. simulans* about 3 million years ago. (right) Schematic of the inter-specific swap of the *trabl* gene by either *D. melanogaster* or *D. simulans* allele using CRISPR/Cas9. (**B**) RNA *in situ* HCR against *Stellate* in testes with *trabl*^*mel*^ or *trabl*^*sim*^. (**C**) RNA *in situ* HCR against *aub* and *AGO3* in testes with *trabl*^*mel*^ or *trabl*^*sim*^. (**D**) Fertility tests of males with *trabl*^*mel*^, *trabl*^*sim*^, and *trabl*^*sim*^ with simultaneous *Stellate* KD, Aub expression, or AGO3 expression (by *nos-* and *bam-*GAL4). (**E**) Scatter plot showing the differential gene expression in *D. simulans* testes of *trabl* mutants and controls. (**F**) Fold-reduction of *aub* and *AGO3* expression upon *trabl* perturbations in *D. melanogaster* and *D. simulans*. (**G**) Schematic showing the three domains of Trabl with functional annotations. MK tests of sliding windows (100 amino acids in size) along Trabl identified a single region within the ZNF domain that shows signature of positive selection. Fisher’s exact test p-value: 0.047. (**H**) Schematic of the inter-specific swap of each of the three domains of Trabl by orthologous *D. simulans* sequences using CRISPR/Cas9. (**I**) RNA *in situ* HCR against *Stellate* in testes with *trabl*^*simZAD*^, *trabl*^*simIDR*^, or *trabl*^*simZNF*^. (**J**) Fertility tests of males with *trabl*^*simZAD*^, *trabl*^*simIDR*^, or *trabl*^*simZNF*^. (**K**) Illustration of the DNA binding innovation in Trailblazer that up-regulated *aub* and *AGO3* expression to boost the genome defense capacity.

To understand the function of Trailblazer in *D. simulans*, we mutated the *trailblazer* ortholog using CRISPR/Cas9 by replacing the entire gene with a DsRed expression cassette. RNA-seq analysis of testis gene expression revealed that, unlike in *D. melanogaster, aub* and *AGO3* expression remained virtually unchanged in *D. simulans trailblazer* mutants (Fig. 4, E and F).

Thus, the ability of Trailblazer to drastically up-regulate *aub* and *AGO3* is a recent innovation in *D. melanogaster* that paralleled the evolution of selfish *Stellate* genes and their *Su(Ste)* repressors.

To further examine the evolutionary changes in the Trailblazer protein, we conducted structure-function analysis of Trailblazer. Trailblazer has a ZAD domain at the N-terminus, an intrinsically disordered region (IDR) in the middle, and an array of five zinc-fingers (ZNF) at the C-terminus (Fig. 4G). In many transcriptional regulators, the ZNF array is responsible for sequence-specific binding to DNA. By fusing different Trailblazer domains to the DNA-binding domain of yeast transcription factor GAL4 and expressing them in *Drosophila* Schneider 2 cells, we found that ZAD and IDR, but not ZNF, domains can activate transcription of the UAS reporter gene (Fig. S3A). In fact, expressing *trailblazer* transgenes that lack either the ZAD or IDR domain in ovarian somatic cells can still activate *aub* and *AGO3* expression (Fig. S3B), suggesting that either domain is sufficient for transcriptional activation but neither is required for DNA binding. Meanwhile, the *trailblazer* lacking the ZNF domain fails to induce *aub* and *AGO3* expression in ovarian soma (Fig. S3B), consistent with the role of ZNF in DNA binding. Therefore, the transcriptional activation and the DNA binding functions can be assigned to the ZAD-IDR and ZNF domains, respectively.

Next, we performed McDonald-Kreitman (MK) tests^36^ to identify signals of positive selection within different domains of Trailblazer (Fig. 4G). This analysis yielded a single region within the ZNF domain that contains an excess of fixed, nonsynonymous substitutions between *D. melanogaster* and *D. simulans* (alpha: 0.85, Fisher’s exact test p-value: 0.047)^37^ (Fig. 4G), suggesting a history of adaptive evolution in the DNA-binding ZNF domain of Trailblazer. To experimentally interrogate the functional impact of evolutionary changes in the Trailblazer protein, we used CRISPR/Cas9 to swap each of the three domains—ZAD, IDR, and ZNF—of the *D. melanogaster* protein with orthologous sequences from *D. simulans* (Fig. 4H). Remarkably, swapping the ZNF, but not ZAD or IDR, domain from *D. simulans* disrupted *Stellate* silencing and compromised male fertility (Fig. 4, I and J), demonstrating that critical evolutionary changes required for *Stellate* silencing occurred in the DNA-binding ZNF domain of Trailblazer. We conclude that recent innovation in the DNA-binding domain of Trailblazer allowed it to up-regulate *aub* and *AGO3* to combat ampliconic *Stellate* genes in *D. melanogaster* (Fig. 4K).

### Escalating defense capacity to tame expanding selfish genes as a conserved strategy

Taken together, we discovered a novel genome defense factor, Trailblazer, that is critical for the control of recently evolved selfish *Stellate* genes in *D. melanogaster*. Trailblazer is a germline transcription factor that promotes the expression of Aub and AGO3, two effectors in the piRNA pathway that silence *Stellate* genes via enzymatic cleavage of their mRNAs. Remarkably, Trailblazer does not regulate Aub and AGO3 expression in testes of closely related *D. simulans* species, which lacks the *Stellate/Su(Ste)* system. Furthermore, while Trailblazer is required for *Stellate* silencing in the male germline of *D*.*melanogaster*, it appears dispensable in the female germline, suggesting that Trailblazer represents defense innovation against male-specific, selfish *Stellate* genes. Importantly, male fertility upon *trailblazer* perturbations is only abolished when the *Stellate* copy number is high, suggesting that Trailblazer reflects a defense adaptation to the *Stellate* copy-number expansion. We propose that, ancestrally, the piRNA pathway was able to tame *Stellate* genes without increasing Aub and AGO3 expression. However, further expansion of the *Stellate* genes ultimately surpassed the silencing capacity of the piRNA pathway, leading to selection for Trailblazer innovations that drive higher levels of Aub/AGO3.

The ability of Trailblazer to markedly promote Aub and AGO3 expression in *D. melanogaster*, but not in *D. simulans*, is driven by recent innovation in its DNA-binding domain, which is accompanied by a 7-fold higher expression level of *AGO3* in *D. melanogaster* than *D. simulans* testes (Fig. 1C). In fact, depletion of *trailblazer* caused a strong reduction in the total piRNA abundance in *D. melanogaster* testes (Fig. 1B). Even more strikingly, the piRNA landscape of *D. melanogaster* testes upon *trailblazer* KD resembles that of wildtype *D. simulans*, lending strong support to the notion that Trailblazer is responsible for the elevated piRNA level in *D. melanogaster* testes compared to *D. simulans* testes.

Our discovery of Trailblazer reveals that genome defense against lineage-specific selfish genes requires genetic innovations beyond specificity factors. In addition to the innovation of *Su(Ste)* piRNAs, which enable the recognition of *Stellate*, Trailblazer innovations that up-regulate Aub and AGO3 expression are paramount, highlighting a quantitative demand for control of ampliconic selfish genes. Copy-number expansion is a recurrent feature of diverse selfish genes^9– 12^, the control of which may require similar tuning of genome defense capacities. For example, a different selfish meiotic driver, *Distorter on the X chromosome* (*Dox*), which jeopardizes *D. simulans* male fertility, arose recently after the split of *D. simulans* and *D. melanogaster* and also underwent a radical copy-number expansion^38–40,10,11^. Previous studies have shown that *Dox* is controlled by the siRNA pathway, instead of the piRNA pathway. In this case, the evolution of *Dox*-targeting siRNAs allowed the siRNA pathway to silence *Dox*—another example of the well-recognized defense strategy in innovating specificity factors to target emerging selfish genes. However, similar to *Stellate, Dox* is ampliconic, so our model would predict a quantitative tuning of the silencing capacity of the siRNA pathway to facilitate *Dox* silencing. Supporting this conjecture, two key components of the siRNA pathway—Dcr-2 (which generates siRNAs) and AGO2 (which cleaves target RNAs identified by siRNAs)—show 2-fold higher expression in *D. simulans* than *D. melanogaster* testes (Fig. 1C). This observation suggests that Dcr-2 and AGO2 might be up-regulated in *D. simulans* to tame expanded selfish *Dox* genes, while AGO3 is up-regulated in *D. melanogaster* to tame expanded selfish *Stellate* genes.

In tetrapods, different retrotransposon families underwent lineage-specific expansions, and many of them are controlled by the expanded KRAB zinc finger protein family. While much work has focused on the innovation of KRAB-ZFPs recognizing individual retrotransposons, our work predicts that controlling the expanding retrotransposons might also involve tuning general silencing machinery, such as KRAB-associated protein-1 (KAP1)—the shared co-factor that mediates the silencing guided by diverse KRAB-ZFPs^41,42^. Even more broadly, we envision that tuning the general defense machinery to quantitatively match the abundance of selfish genes is a widespread phenomenon and a central strategy in evolutionary battles against the expanding selfish genes across the tree of life.

## Acknowledgments

We thank Andy Clark, Bruce Hay, Ching-Ho Chang, Ildar Gainetdinov, and Lu Yue for comments on the manuscript. We thank Ellen Rothenberg for discussion. We thank James McGehee and David Stern for advice on CRISPR/Cas9 in *D. melanogaster* and *D. simulans*, respectively. We thank Toshie Kai, Mayu Inaba, Bloomington Drosophila Stock Center, Vienna Drosophila Resource Center, and Kyoto Drosophila Stock Center for fly stocks. We are grateful to Elena Udartseva and Baira Godneeva for technical assistance. We appreciate the help of Igor Antoshechkin with sequencing, the help of Grace Shin with HCR, and the help of Giada Spigolon and Andres Collazo with microscopy.

## Funding

National Institutes of Health, R01GM097363 (AAA)

Howard Hughes Medical Institute, Faculty Scholar Award (AAA)

National Institutes of Health, R35GM142494 (YCGL)

## Author contributions

Conceptualization: PC

Methodology: PC

Investigation: PC, KCP, EHP, YL, YCGL

Visualization: PC

Funding acquisition: AAA, YCGL

Project administration: PC, AAA

Supervision: PC, AAA

Writing – original draft: PC

Writing – review & editing: PC, KCP, EHP, YL, YCGL, AAA

## Methods

### Fly husbandry and strains used

Flies were raised at 25°C and 70% relative humidity. All fly lines used in this work were listed in Table S1 (for the screen) and Table S2 (for all other experiments). For each panel in main figures, all flies have the same X chromosome and thus identical *Stellate* copy-numbers. For all KD experiments (including the RNAi screen), a double-driver line carrying both *nos*-GAL4 (on chr3) and *bam-*GAL4 (on chrX) was used, and it was always used as the mother, such that all male progeny inherit the same X chromosome carrying *bam-*GAL4 and the same *Stellate* copy-number. The genotypes and the X chromosome status (i.e., *Stellate* copy number) of each main figure panel are listed in Table S3.

### Transgenic fly (*D. melanogaster*)

To drive *trabl* expression in flies, we made *UASp-CG17801* transgenes with a GFP tag at either the N- or the C-terminus, followed by K10 3’UTR. The region targeted by the RNAi line (TRiP.HMC05112, BDSC # 60118) was recoded to synonymous sites to evade the KD (from <u>ACa TCg ATt AAg tTa AAg</u> to <u>ACg TCc ATc AAa cTg AAa</u>). To drive expression of truncated *trabl* transgenes, each of the three domains was deleted from the *UASp-CG17801-GFP* construct: ΔZAD (Δaa8-80), ΔIDR (Δaa81-229), and ΔZNF (Δaa230-368). The transgene along with a mini white marker was integrated at either the VK37 or VK31 attP site on chr2 and chr3, respectively, by PhiC31 integrase. To swap the endogenous *trailblazer* gene by intended sequences, we employed CRISPR/Cas9. Two guide RNAs flanking the *trailblazer* gene were selected using the web tool Target Finder^43^ (<u>targetfinder.flycrispr.neuro.brown.edu/</u>)—GTT TCA TGG CGT TAC CAG AT <u>AGG</u> and CAT AAC GAC CCT TGA ATA AA <u>AGG</u> (underlined are the PAM sites)—and cloned into the pCFD5 plasmid^44^. For the homology-directed repair template, we adapted a 3xP3-DsRed marker (flanked by piggyBac termini) from the pScarlessHD plasmid (a gift from Kate O’Connor-Giles, Addgene plasmid # 64703) and introduced two ∼1kb homology arms flanking the two guide RNA target sites. The *trailblazer* coding region was replaced by 1) *trabl*[mel] cDNA, 2) *trabl*[sim] cDNA (XM_002102980), 3) *trabl*[simZAD] cDNA (aa1-80), 4) *trabl*[simIDR] cDNA (aa81-243), or 5) *trabl*[simZNF] cDNA (aa244-396). Once the successful editing was obtained, the DsRed marker was removed by introducing the PiggyBac transposase. All injections, screenings, and balancing were done by BestGene (Chino Hills, CA).

### Transgenic fly (*D. simulans*)

To delete the *trailblazer* ortholog in *D. simulans*, we used CRISPR/Cas9 to replace the entire gene by 3xP3-DsRed, employing the homology-directed repair as done in *D. melanogaster*. The two guide RNA sequences are: CAG CAA AAT TAG CTG TTA AA and TAA TCC CGT CTT GGA AGA TG. To increase the editing efficiency, the guide RNA expression unit and the homology-directed repair template were combined into one plasmid^45^. The plasmid was injected into a *nos-Cas9* strain^46^ by Genetivision (Houston, TX). The successful editing was first shortlisted by DsRed expression in the eye and then confirmed by genomic PCR.

### Fertility tests

Virgin flies were allowed to individually mate with two wildtype (*yw*) virgin flies of the opposite sex for 3 days. They were then flipped to a new vial, where they mated for another 3 days. This was repeated one more time before flies were discarded, resulting in a total of 3 vials and 9 days of mating per tested fly. Offspring in a given vial was counted 17 days after the initial introduction of flies, so eggs in each vial had 14-17 days to develop to adulthood. Three counts from the 3 vials were summed to produce the total progeny of a tested fly, which is shown as a single dot in the fertility test plots in this work.

### RNA *in situ* HCR, RNA FISH, and immuno-fluorescence

Testes (3-7 days old) or ovaries (3-7 days old, yeast-fed for 2 days) were dissected in PBS and fixed in 4% formaldehyde in PBS with 0.1% Tween 20 (PBSTw) for 20mins. Gonads were then washed in PBSTw three times before being permeabilized in PBS with 0.5% Triton X-100 (PBST) for 30mins. For RNA *in situ* HCR, downstream procedure was done according to manufacturer’s recommendations for generic samples in solution. Buffers, probes, and amplifiers for HCR v3 were purchased from Molecular Instruments. HCR targets and the unique identifiers of respective probe sets are: *Stellate* (4537/E832), *trabl* (5015/F460), *aub* (5040/F508), and *AGO3* (5040/F510). For RNA FISH against *Su(Ste)*, DNA probes against position 994-1068 of *Su(Ste): CR42424* with fluorophore conjugation were purchased from IDT. RNA FISH was done following the HCR protocol but omitted the signal amplification step. For immuno-fluorescence, gonads were blocked in PBSTw with 5% BSA for 1hr after fixation and permeabilization. Next, gonads were incubated in PBSTw with primary antibody against Fas3 (DSHB 7G10, 1:200) at 4°C overnight, washed three times in PBSTw, incubated in PBSTw with secondary antibody for 1hr at room temperature, and washed three times in PBSTw. Finally, gonads were mounted in VECTASHIELD Medium with DAPI (H-2000). To compare mature sperm in seminar vesicles, unmated virgin males were aged for 7 days before dissection. For all other comparisons, animals of the same age were used.

### Imaging acquisition and processing

Confocal images were acquired with Zeiss LSM 980 using a 63x oil immersion objective with 1.4 NA and processed using Fiji. Within each figure panel, identical settings were used across samples at both the image acquisition and processing steps. All images shown are from single focal planes. Dotted outlines of testes were manually drawn based on the DAPI channel for illustration purposes, and asterisks were used to mark the hub (a cluster of somatic cells that serve as the stem cell niche) at the apical tips of testes.

### RNA-seq and analysis

RNA-seq was done as described in our recent work^24^. Briefly, testes from *D. melanogaster* males of *trabl* or *gfp* germline KD (driven by *nos-* and *bam-*GAL4) and testes from *D. simulans* males of *trabl* heterozygotes or *trabl* mutants were dissected (20-30 testes per sample) and RNA was extracted using TRIzol. Polyadenylated RNA was selected using NEBNext Poly(A) mRNA Magnetic Isolation Module (E7490S), library prepped using NEBNext Ultra II Directional RNA Library Prep Kit (E7760S), and sequenced on NextSeq2000. For *D. melanogaster*, transcripts from two biological replicates per genotype were quantified by kallisto^47^, using an index file that combines the transcriptome and repeat sequences from RepBase. Paralogs of *Stellate* genes were merged to show overall expression of the *Stellate* gene family. Differential gene expression was analyzed using DESeq2^48^. For *D. simulans*, in addition to sequenced samples, wildtype (*w*^*501*^) testis RNA-seq data of two biological replicates were downloaded from NCBI SRA (SRR24223890 and SRR24223891)^38^, and the transcriptome associated with the assembly GCF_016746395.2 was downloaded from NCBI for kallisto quantification.

### Small RNA-seq and analysis

Small RNA-seq was done as described in our recent work^30^. Briefly, testes from males of desirable genotypes were dissected in PBS. The Argonaute-associated small RNAs were isolated from testes using the TraPR columns^49^, library prepped using NEBNext Small RNA Library Prep Set (E7330S), and sequenced on NextSeq2000. The adaptor sequences were trimmed using cutadapt. For *D. melanogaster*, two genotypes were analyzed: *trabl* and control (*gfp*) germline KD (driven by *nos-* and *bam-*GAL4). For *D. simulans*, the *w*^*501*^ wildtype strain was used.

### ChIP-seq and analysis

ChIP-seq was done as described in our recent work^24^. Briefly, ovaries (*nos*- and *bam-*GAL4 > UASp-Trabl-GFP) from 50 yeast-fed females were dissected in PBS and then fixed in 1% formaldehyde for 10mins. After three PBS washes, fixed ovaries were snap-frozen in liquid nitrogen and stored at -80°C until ChIP experiments. Frozen ovaries were thawed in Farnham Buffer and then homogenized in RIPA buffer using a glass douncer and a tight pestle. Next, samples were sonicated in Bioruptor (Diagenode) on high power for 25 cycles (30s on and 30s off) and centrifuged to obtain the supernatant. The supernatant was pre-cleared with Dynabeads Protein G beads for 2hrs, then with 5% being set aside as the Input and the rest being incubated with 5μl anti-GFP antibody (A-11122, Invitrogen) overnight at 4°C. The immuno-precipitated (IP) sample was incubated with Dynabeads Protein G beads for 5hrs at 4°C, followed by five LiCl washes. At the same time, the Input sample was incubated with RNase A for 1hr at 37°C. Finally, both IP and Input samples were incubated with protein K at 55 □C for 3hrs and then at 65 □C overnight (with intermittent shaking). DNA was isolated by phenol-chloroform extraction and quantified by Qubit. A total of two biological replicates of ChIP were done, and they were then subject to library preparation using NEBNext Ultra II DNA Library Prep (E7103S) and sequenced on NextSeq2000. Reads were mapped to the dm6 genome assembly using bowtie, and peaks were called using macs2 callpeak with the default setting. IGV (igv.org) was used to generate genome browser tracks.

### Estimation of the *Stellate* copy number

To estimate the *Stellate* copy number on a given X chromosome, the line of interest was first crossed to C(1)RM to generate XO males that lack Y chromosomes, as Y-linked *Su(Ste)* sequences are homologous to X-linked *Stellate* sequences and could confound the *Stellate* copy-number estimation. To extract genomic DNA of XO males bearing the X chromosome from the line of interest, XO males were homogenized, incubated with proteinase K for 1hr at 55°C, incubated with RNase A for another 1hr at 37°C, and then subject to phenol-chloroform extraction and ethanol precipitation. Quantitative PCR was done on extracted genomic DNA targeting a single-copy gene (*zuc*) and *Stellate*, and the copy number of *Stellate* was estimated by the abundance of *Stellate* DNA normalized to that of *zuc*. The primer sequences are TTT GGC GGA TTC AAT AAA GC (*zuc* forward), TTG TGC ATC AAG TTC GTG GT (*zuc* reverse), ATC GAT TGG TTC CTC GGG ATC AA (*Stellate* forward), and GCC GTA CAA CAA GCC AGA GGA ACT A (*Stellate* reverse). The lines surveyed are listed in Table S3.

### Mapping the trans-activation domain

*Drosophila* Schneider 2 (S2) cells were grown at 25°C until they achieved a confluency range of approximately 70% to 80%. The cells were then subjected to co-transfection of a UASp-GFP-NLS reporter plasmid and another plasmid expressing GAL4 DNA-binding domain (DBD) fused to different domains—1) GAL4 trans-activation domain (as a positive control), 2) nothing (as a negative control), 3) Trabl ZAD, 4) Trabl IDR, or 5) Trabl ZNF—under the *actin 5C* promoter. Transfection was done for three replicates, using the TransIT-LT1 Transfection Reagent (Mirus Bio MIR2304) following the manufacturer’s instructions. Transfected cells were harvested two days later, and RNA was isolated using TRIzol. Reverse transcription was done by SuperScript III (Invitrogen) with random hexamers, and the mRNA expression levels of GFP and GAL4 DBD were measured by quantitative PCR. The primer sequences are TAC AAC AGC CAC AAG GTC TAT ATC A (*gfp* forward), GGT GTT CTG CTG GTA GTG GTC (*gfp* reverse), AGT GTC TGA AGA ACA ACT GGG AG (GAL4 DBD forward), and GCT GTT CCA GTC TTT CTA GCC TT (GAL4 DBD reverse). The trans-activation domain of Trabl is expected to activate transcription after being recruited to UASp sites by GAL4 DBD.

### McDonald-Kreitman test

McDonald-Kreitman (MK) test, which contrasts the ratio of nonsynonymous and synonymous divergence to that of polymorphism^36^, was conducted using polymorphism within a *D. melanogaster* Zambian population and divergence between *D. melanogaster* and *D. simulans* (XP_002103016.2). The number of nonsynonymous and synonymous changes was counted using the mutational paths that minimize the number of amino acid changes, and the significance of the MK test using Fisher’s exact test. Sliding window MK tests were performed with windows of 100 codons and 10-codon steps.

**Fig. S1.**
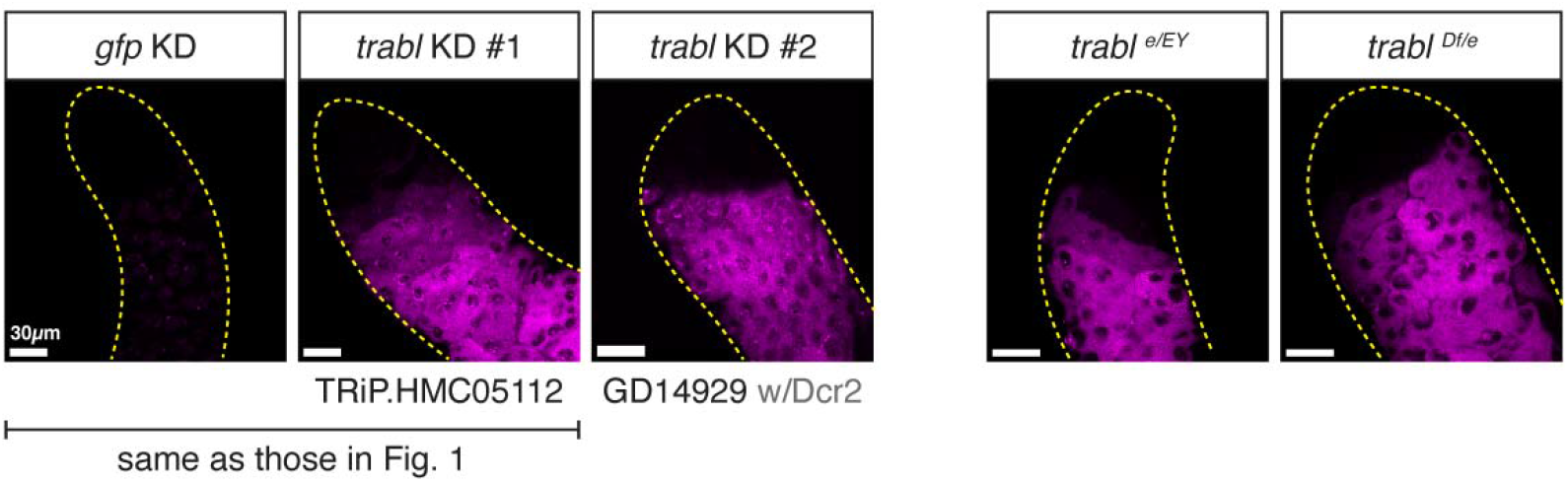
RNA in situ HCR against Stellate in testes upon trabl perturbations by knock-down or mutations. The RNAi lines and mutant alleles used are indicated. UAS-Dcr-2 was included for the GD line.

**Fig. S2.**
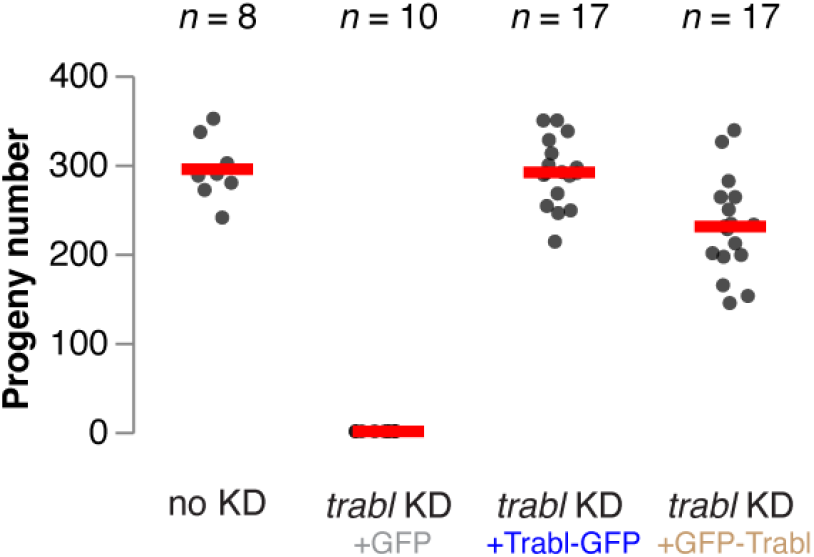
Male fertility tests upon no KD and trabl germline KD with simultaneous GFP, Trabl-GFP, or GFP-Trabl expression driven by nos- and bam-GAL4.

**Fig. S3.**
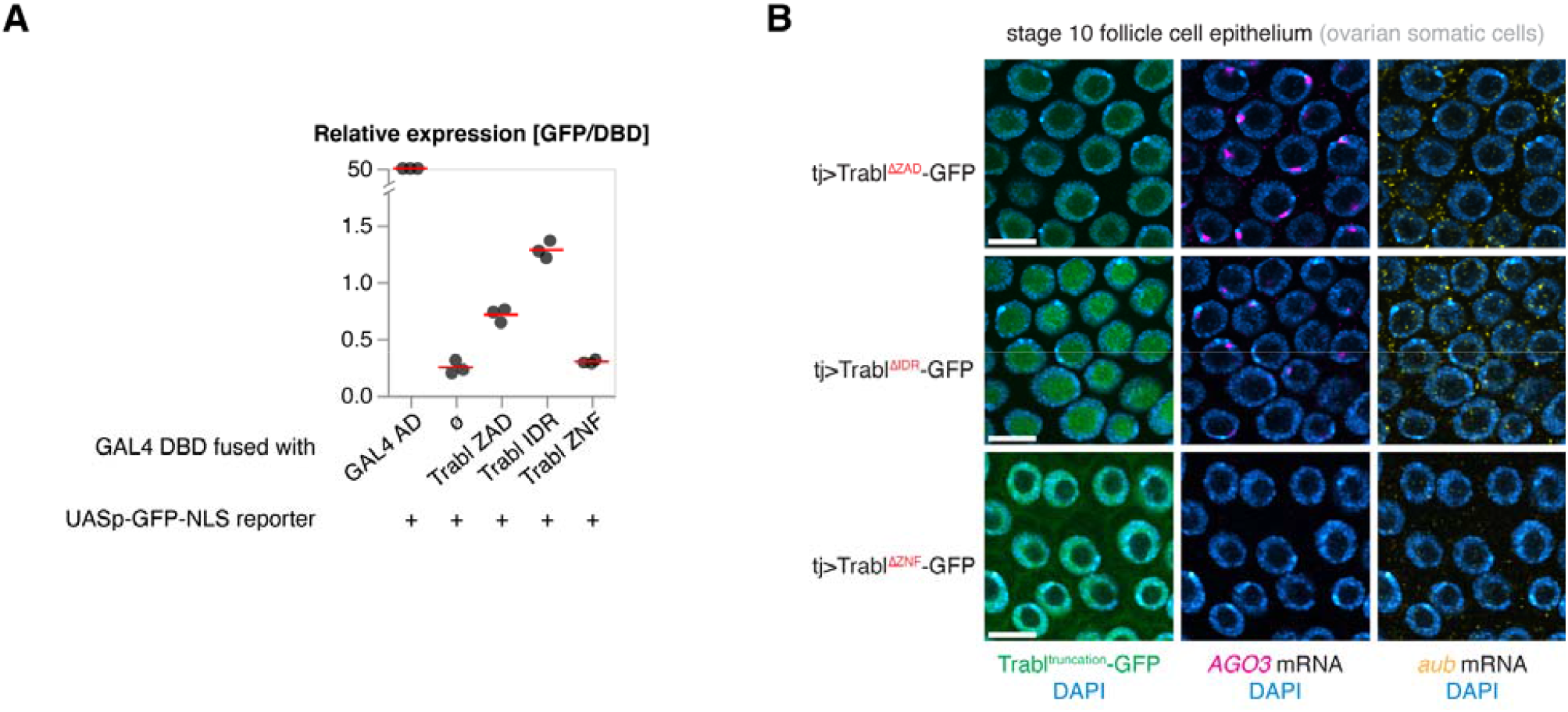
(**A**) Relative expression of GFP (normalized to DBD) quantified by RT-qPCR upon co-transfection of UASp-GFP-NLS and another plasmid expressing GAL4 DBD fused to 1) GAL4 AD, 2) nothing, 3) Trabl ZAD, 4) Trabl IDR, or 5) Trabl ZNF. Red lines mark the average of three replicates. (**B**) RNA *in situ* HCR against *aub* and *AGO3* in follicle cell epithelium with forced expression of Trabl[ΔZAD]-GFP, Trabl[ΔIDR]-GFP, or Trabl[ΔZNF]-GFP transgenes by *tj*-GAL4. Note that nucleoli appear as DAPI-weak circles inside the follicle cell nuclei.

